# Modeling the *C*. *elegans* Germline Stem Cell Genetic Network using Automated Reasoning

**DOI:** 10.1101/2021.08.08.455525

**Authors:** Ani Amar, E. Jane Albert Hubbard, Hillel Kugler

## Abstract

Computational methods and tools are a powerful complementary approach to experimental work for studying regulatory interactions in living cells and systems. We demonstrate the use of formal reasoning methods as applied to the *Caenorhabditis elegans* germ line, which is an accessible model system for stem cell research. The dynamics of the underlying genetic networks and their potential regulatory interactions are key for understanding mechanisms that control cellular decision-making between stem cells and differentiation. We model the “stem cell fate” versus entry into the “meiotic development” pathway decision circuit in the young adult germ line based on an extensive study of published experimental data and known/hypothesized genetic interactions. We apply a formal reasoning framework to derive predictive networks for control of differentiation. Using this approach we simultaneously specify many possible scenarios and experiments together with potential genetic interactions, and synthesize genetic networks consistent with all encoded experimental observations. *In silico* analysis of knock-down and overexpression experiments within our model recapitulate published phenotypes of mutant animals and can be applied to make predictions on cellular decision-making. This work lays a foundation for developing realistic whole tissue models of the *C*. *elegans* germ line where each cell in the model will execute a synthesized genetic network.

## 1. Introduction

Understanding the mechanisms and design principles underlying the transition of stem cells from self-renewal to differentiation is a fundamental question in biology. The control of these self-renewal and differentiation processes is crucial for proper organ formation, for tissue maintenance and repair, and has important implications in unravelling the molecular and cellular basis of tumor initiation and progression [1, 2]. The *Caenorhabditis elegans* germ line provides an accessible model system for studying stem cell decision-making. Over the last decades, a number of regulatory interactions between core signaling genes governing germ cell identity has been experimentally determined (see [3] and references within). The current understanding of the decision of cells to remain stem cells or to differentiate involves the interaction of multiple pathways that are inhibited by active Notch pathway signaling. The genetic interactions between these pathways and their components are complex. Computational modeling, especially cell-based modeling, can play an important role in uncovering proposed interactions that are inconsistent with available experimental data or those that may be most informative in clarifying the network structure and dynamics and therefore deserve more experimental focus.

Several computational models for studying the *C*. *elegans* germ line have been developed in the past years (see [4] and references within), particularly in the context of proliferative germ cell behavior. Among the first models, exploring germ line development (larva) and maintenance (adult), is the model proposed by Setty [5]. Although the computational approach made simplifying assumptions regarding the spatial aspects of the model (assuming a 2D geometry and a discrete grid in which cells can be positioned), this model nevertheless successfully represented germ cell movements inside the gonad, including the interplay between an individual cell’s cycle control and differentiation in response to changes in signaling. The limitations in describing spatial aspects in [5] were overcome in a mechano-logical model of the germ line [6], which utilized a more realistic 3D simulation of germ cell movement and distal tip cell (DTC) migration. In both models, germ cell behavior was represented by the statecharts approach [7, 8], which extends classical state machines with orthogonal states, a hierarchy of states and an executable language enabling efficient code generation and simulation. Orthogonal states are used for defining various aspects of cellular behavior (e.g., cell cycle, signaling genes, cell fate decision), using transitions between states for each component (within the object), and the conditions required for each transition. There are some important differences between the two models. First, the Atwell model [6] has a more realistic representation of cell spatial location and direction of movement (without the need to be uniformly spaced on a fixed grid). However, this off-lattice approach to cell mechanics and special algorithms to efficiently handle cell-cell neighborhoods increased the computational cost, in comparison to the lattice-based model of Setty. Second, the Atwell model was applied to more accurately simulate cell tracking and labeling throughout the process of gonad development. This is especially useful for studying the interrelationship between the gonad and the germ line, and between cells within the germ line over time – a challenging aspect of experimental approaches.

Here we present a first detailed network model for germline stem cells, that explores the specification of the cell fate in *C*. *elegans* by means of state-of-the-art formal reasoning synthesis methods, and the reasoning engine for interaction networks tool (RE:IN) [9, 10, 11]. RE:IN is a synthesis-based tool, that is now available as an open source data science framework (the reasoning framework) that supports scalable formal reasoning procedures combined with a user friendly interface to specify interaction network models constrained by experimental results. Synthesis approaches for biological modeling are becoming an important area of research and applications, see for example [12, 13, 14, 15, 16, 17, 18, 19, 20] and references within. A significant advantage of synthesis compared to our earlier work [5, 6] is the automation and computational efficiency enabled by the formal reasoning approach, providing the identification of networks that satisfy a wide set of biological constraints specified by experimental results. This prevents the bias introduced by manually searching for one or few networks that are plausible in a trial and error process. We computationally synthesize networks that capture signaling and genetic interactions within a single germ cell’s decision-making process, focusing on the transition between the undifferentiated stem/progenitor fate and differentiation (entry into the meiotic pathway). Such synthesized networks can ultimately be incorporated within computational agents representing cells to study the development of the entire germ line [5, 6], resulting in models that can be simulated and compared to in vivo phenotypes.

Based on published data and known genetic interactions, we constructed a genetic network, which represents the “stem cell fate versus meiotic development” decision circuit in the young adult hermaphrodite germline stem cell system and which extends previous models in a number of ways. First, our network model is more detailed in terms of intracellular signaling (number of components and interactions) and is derived from extensive studies from the literature. Second, we applied the automated approach for constructing and refining a suitable model that guarantees satisfying the set of all specified biological constraints and experiments. Third, the presented approach allows the synthesis algorithm to find all existing networks (by selecting a subset of the optional interactions) consistent with encoded experimental behavior, unlike other approaches that account for one or a limited set of scenarios.

The model enables us to explore the behavior within an individual cell, that either resides in a niche in the range of the DTC signal or just out of range of the DTC signal, or that loses DTC signaling (when moving from distal to proximal part of the germ line). We analyzed the structure and dynamics of this genetic network, its functional modules and derived predictive networks for control of self-renewal and differentiation. *In silico* analysis of knock-down and over expression experiments within our model recapitulate published phenotypes of mutant animals and is applied to make new predictions on cellular decision making.

Below is a summary of the steps we have found important for the development and analysis of this model:

1. Extensive study of published experimental data and known /hypothesized genetic interactions in the *C*. *elegans* germ line.
2. Preparation of data collection table that determines “source” (starting node) and “target” (ending node) components, the interactions between them and data source (see Table 1 for an example).

**Table 1.** Regulations of genes as depicted in Figure 1 and their literature references.
3. Development of logical rules leading to an activity of each component, using the Boolean operators AND, OR, and NOT.
4. Encoding a set of components and interactions in the reasoning framework.
5. Visualisation of the developed network in the reasoning framework and its refinement against data collection table and logical rules.
6. Encoding a set of experimental observations that incorporates all of the information. The corresponding network model files are provided in Supplementary data.
7. Selection of analysis options (e.g., experiment length, update scheme) and running the solver.
8. Assessment of the found solutions (if any) vs. expected profiles of gene activity.
9. Revision of model assumptions (see steps 2, 3 and 6) if no model solutions exist.
10. Once confirmed to be consistent with experimental data, the model was extended by adding *in silico* genetic perturbations (known phenotypes) to the set of experimental observations.
11. Comparison of the found solutions to the results of known phenotypes.
12. Formulation and evaluation of new predictions that have not yet been experimentally observed.

| Source | Regulates | Target | Reference |
| --- | --- | --- | --- |
| DTC | act | APX1 | [23] |
| DTC | act | LAG2 | [24] |
| LAG2 | act | GLP1 | [24] |
| APX1 | act | GLP1 | [23] |
| GLP1 | act | GLP1ICD | [25] |
| GLP1ICD | act | SEL8 | [26] |
| GLP1ICD | act | LAG1 | [26] |
| SEL8 | act | LAG1 | [26] |
| GLP1 | act | HOP1 | [27] |
| HOP1 | act | GLP1ICD | [27] |
| GLP1 | act | SEL12 | [28] |
| SEL12 | act | GLP1ICD | [28] |
| SEL10 | rep | GLP1ICD | [29] |
| SEL10 | rep | SEL12 | [30] |
| LAG1 | act | LST1 | [31] |
| LAG1 | act | SYGL1 | [31] |
| LST1 | rep | GLD1 | [22] |
| LST1 | rep | FBF1 | [22] |
| FBF1 | rep | GLD1 | [22] |
| SYGL1 | rep | FBF1 | [22] |
| LAG1 | act | FBF2 | [32] |
| FBF2 | rep | GLD1 | [22] |
| FBF | act | GLD3 | [33, 34] |
| FBF1 and FBF2 | rep | GLD3 | [35] |
| GLD3 | act | GLD2 | [35] |
| NOS3 | act | GLD1 | [36] |
| GLD2 | act | GLD1 | [36] |
| FBF1 | rep | SCF<sup>PROM-1</sup> | [37] |
| FBF1 and FBF2 | rep | AxisandSCprotein | [38] |
| SCF<sup>PROM-1</sup> | rep | CYE1/CDK2 | [39] |
| "act" for | activate; "rep" for repress |  |  |

## 2. The Reasoning Framework

To investigate the dynamics of this genetic network and potential regulatory interactions, we used the reasoning framework (for more information see Yordanov et al.[11] and the Materials and Methods section). This approach supports the modeling of gene networks via Abstract Boolean Networks (ABN). In this logical modeling framework, an ABN contains a set of components (e.g., genes / signals / transcription factors / proteins) which can be active or inactive (represented by a Boolean value) and interactions (definite or possible) which can be either positive (activation) or negative (inhibition). In addition, a set of (18) regulation conditions allows choice in defining how interactions are combined to determine the effect of multiple interactions on a given component. For example, if two components g1 and g2 regulate component g, the choice of the regulation condition will determine if both g1 and g2 are required to activate g (AND logical function) or either g1 or g2 are sufficient to activate g (OR logical function). In general, the current 18 regulation conditions supported in the reasoning framework take into account multiple regulators (activators and inhibitors), and define the activation of a gene as a logical function depending on the activity state (active/inactive) of these regulating components (see [11] and Supplementary Material, Figure S1). The regulation conditions distinguish between a case where all, some or none of the activators (inhibitors) are active and have a property of monotonicity, implying that if a regulated component is active, and one of its activators switches from inactive to active, the regulated component will remain active, and similarly if a regulated component is inactive, and one of its inhibitors switches from inactive to active, the regulated component will remain inactive. There are several features of this approach that should be emphasized in terms of the current genetic network construction:

1. The formal reasoning-based synthesis algorithms are utilized to efficiently explore the large state space of potential networks and to find solutions that recapitulate all specified experimental observations.
2. Each consistent network is a concrete Boolean network, which includes a specific subset of the *optional* interactions, all the *definite* interactions, and one regulation condition for each component. For example, a target gene may requires the following regulation condition (regulation condition 5, Supplementary Material, Figure S1):

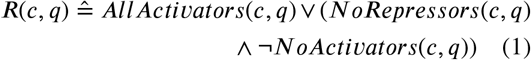

where *AllActivators(c,q)* indicates that all activators of a component *c* are present in a state *q*, *NoRepressors(c,q)* indicates that no repressors of *c* are present in *q*. And, by negating the function *NoActivators(c,q)*, we obtain the function *(¬NoActivators(c,q))* that indicates that some but not all activators of *c* are present in state *q*. In this case, the solver can select which interaction (one of the redundant genes or both of them) and corresponding pathway satisfies the encoded experimental observations.
3. A set of experimental observations (constraints) that a network must satisfy are encoded as predicates over system states, limiting the feasible choices of *optional* interactions and conditions that yield consistent models. For example, in our network, we formalized an observation where the DTC signal is considered to be initially active, but should be inactive at time step 20 (see example in Supplementary Material, Figure S6), simulating a cell in the proximal part of the germ line. In this case, the solver identifies the existing trajectory (if any) that satisfies the following expression:

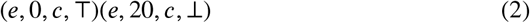

where *e* is the experiment label, *c* denotes the DTC component, T (true) and ⊥ (false).
4. The solver allows the user to set analysis options (e.g., solutions limit, synchronous /asynchronous updates, experiment length – number of time steps).
5. Once the tool finds consistent networks, these networks can be utilized to investigate the network under conditions that were not explicitly specified in the original modelbuilding, and can therefore be used to make prediction of the outcome of new experiments including simultaneous mutations.

## 3. Materials and Methods

We briefly summarize the main formal definitions and concepts underlying the reasoning framework [11, 51, 53]. At the core of the approach are Abstract Boolean Networks, an extension of *Boolean Networks* (BNs) [50, 52]. BNs are a class of GRN models that are Boolean abstractions of genetic systems, i.e., every gene is represented by a Boolean variable specifying whether the gene is active or inactive. The concept of *Abstract Boolean Networks* (ABNs) [9] was introduced to allow the representation of models with network topologies and dynamics that are initially unknown or uncertain. The main goal of the framework is to automatically identify a choice of allowed network topology and dynamics that is consistent with all specified experimental constraints or to prove that no consistent network exists.

Let *G* be a finite set of genes. Let 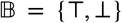 be the Boolean domain and let 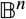 be all vectors of *n* Booleans. Let *E* be a set of directed signed edges between elements of *G*, i.e., 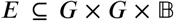. The sign of an interaction is either T for positive regulation (activation) or ⊥ for negative regulation (repression). Let *g* and *g*′ be genes from *G*. We call *g* an *activator* of *g*′ iff (*g*, *g*′, T) ∈ *E*, and a *repressor* iff (*g*, *g*′, ⊥) ∈ *E*. We define the state space of a system as 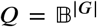. For a given state *q* ∈ *Q* and a gene *g* ∈ *G*, we denote by *q*(*g*) the state of *g* in *q*.

Each gene is associated with an update function *f_g_* with a signature 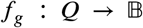 defining its dynamics. For *synchronous* updates, the dynamics of the system are defined in terms of the update functions of all genes applied at each transition, where, given a current state *q* and next state *q*′, ⋀_*g*∈*G*_*q*′(*g*) = *f_g_*(*q*). In this work we focus on synchronous semantics, but asynchronous semantics are also supported by the reasoning framework.

A set of 18 biologically plausible update function templates, called *regulation conditions* are used to determine the dynamics of a gene (see Supplementary Material, Figure S1). Introducing these function ‘templates’ aims to reduce the number of Boolean functions that need to be considered (thus simplifying analysis) while still maintaining and emphasising biological and experimental plausibility.

To capture possible uncertainty and partial knowledge of the precise network topology, we allow some interactions to be marked as *optional* (denoted by the set *E*^?^), each of which could be included in a synthesised *concrete model*. Thus, in terms of network topology, this means a set of 2^|*E*?|^ concrete models, each of which corresponds to a unique selection of possible interactions. Additionally, a choice of several possible regulation conditions for each gene is taken into consideration, leading to the following definition:

An abstract Boolean network is a tuple ⟨*G*, *E*, *E*^?^, *R*⟩, where *G* is a finite set of genes, 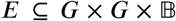 is a set of definite (positive and negative) and directed interactions between them, 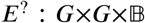 is a set of optional interactions and *R* = {*R*_g_ |⩝_g_ ∈ *G*}, where *R*_g_ specifies a (non-empty) set of admissible regulation conditions for gene *g* [9, 11, 51].

An ABN is transformed into a concrete Boolean network by selecting a subset of the possible interactions to be included and assigning a specific regulation condition to each gene. The semantics of such a concrete model is defined in terms of a transition system 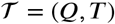, where 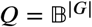 is the state space and *T* is a transition relation defined in terms of the predicate 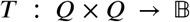. The semantics of the synchronous transition system is then given by

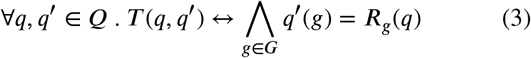

A finite *trajectory* of length *k* is defined as a sequence of states *q*_0_,*q*_1_…,*q*_k-1_ where ⋀_0<*i*<*k*_ · *q*_i_ ∈ *Q* ∧ *T*(q_i-1_,*q*_i_).

A set of *experimental observations* that a BN must be able to satisfy are encoded as predicates over system states, which limits the feasible choices of possible interactions and regulation conditions yielding consistent models. The approach developed and described in [9, 11] allows GRN synthesis: given an ABN and a set of experiments, find a choice of interactions and regulation conditions that guarantees that the resulting concrete BN is consistent with all experimental observations. The synthesis algorithm constructs concrete, consistent models if they exist, or formally proves no solution exists. The reasoning framework is publicly available at (https://github.com/fsprojects/ReasoningEngine).

## 4. Results

### 4.1. Construction of the Model

To construct a genetic regulatory network, we collected literature data about known interactions between the core Notch pathway genes and their downstream effectors regulating stem cell fate versus meiotic development decision in young adult hermaphrodite germline stem cell system (see [3] and references within). Genes were selected for implementation of the circuit as a Boolean model using the reasoning framework [11]. The proposed genetic network includes twenty-four (24) components (genes/gene products) and forty (40) interactions (Figure 1) with a large state space of possible combinations (2^24^ = 16,777,216 states). We defined the network model to be updated under a synchronous update scheme (i.e., the states of all components are updated simultaneously at each step). Interactions that are well characterized and have strong experimental support we denoted as *definite* interactions (represented as solid edges), while interactions that are hypothesized but their current experimental support is weaker or that may be redundant we denoted as *optional* interactions. To reproduce expected profiles of the core *gene activity* (the attributes of gene function that are inferred from genetic manipulation, but that can be the result of changes in levels or function of the resultant mRNA or protein), a set of experimental observations was encoded, which contain input as well as output state of the genes. Furthermore, functional effects of the selected genes were included by encoding knock-out and/or overexpression experimental observations based on known phenotypes (Table 2). To simulate genetic perturbations, a gene was defined as knocked out (KO), or as constitutively active/“overexpressed” (FE = forced expression) along with corresponding experimental constraint (e.g., ‘KO(gene)=1’ or ‘FE(gene)=1’) (for an example of encoding such constraints see Supplementary Files). Once the set of interactions and observations were encoded and the analysis started, the solver searched for all existing networks (if any) that were consistent with all encoded experimental observations. Each network is visualized by presenting the topology of a single specific solution out of many possible solutions (see example in Supplementary Material, Figure S2) and a table representing a simulation of each experiment showing Boolean values of each gene in this solution (see Section 3.3).

**Figure 1:**
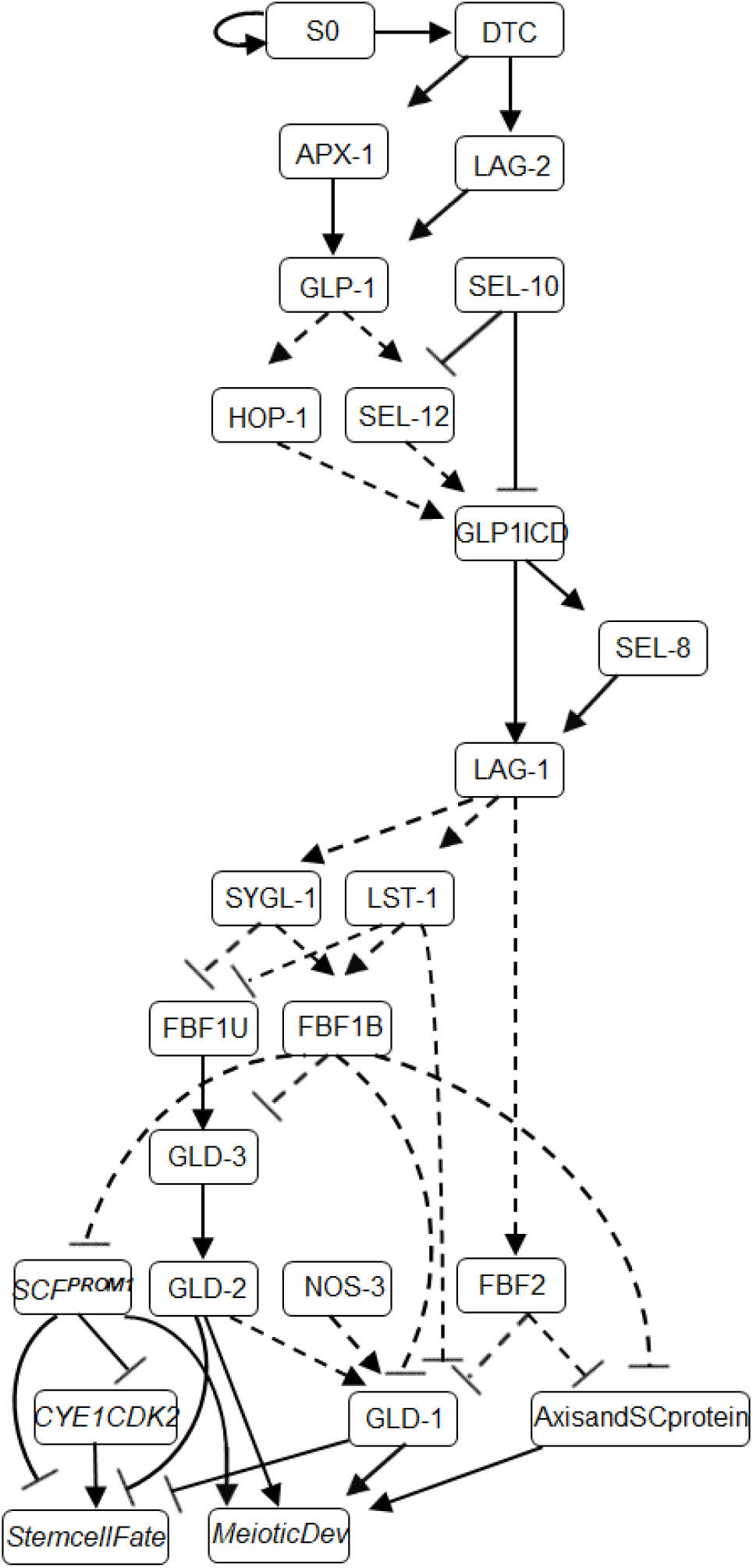
A genetic network model of stem cell fate versus meiotic development. Definite interactions appear as solid lines, optional interactions as dashed lines, arrowhead line corresponds to activation and T-shaped line – to inhibition.

**Table 2.** *In silico* knock-out (KO) and “overexpression” (FE) experiments.

|  | Experiment | Comments | Ref |
| --- | --- | --- | --- |
| KO | GLP-1 | Meiotic entry pathway | [40] |
|  | LST-1 | No effect on stem cell fate | [31] |
|  | SYGL-1 | No effect on stem cell fate | [31] |
|  | LST-1 and SYGL-1 | Meiotic entry pathway | [31] |
|  | GLD-1 and GLD-2 | Failure to enter meiosis | [41] |
| FE | GLP-1 | Failure to enter meiosis | [42] |
|  | LST-1 | Failure to enter meiosis | [34] |
|  | SYGL-1 | Failure to enter meiosis | [34] |

### 4.2. Functional Modules

First, we identified the functional modules (genetic pathways) in the network (Figure 1), associated with specific processes: genes that act downstream of GLP-1 signaling (LST-1, SYGL-1 and FBF-1); the GLD-1, GLD-2 and SCF^PROM-1^ meiotic entry pathways; the GLD-1 accumulation module. It is assumed that the initiation of network interactions occurs due to contact between undifferentiated GLP-1-expressing germ cells and the distal tip cell (DTC), allowing GLP-1 on the surface of the germ cells to bind the membrane-bound ligands LAG-2 and/or APX-1 and to start downstream signaling (for details, see Figure 1 and Table 1). Consequently, DTC – germ cell signaling promotes the stem cell fate by inhibiting the meiotic entry pathways (GLD-1, GLD-2, and SCF^PROM-1^)

Next, we analyzed the kinetics of regulatory module downstream of GLP-1 signaling (LST-1, SYGL-1 and FBF- 1). LST-1 and SYGL-1 proteins are direct transcriptional targets of GLP-1 signaling, that function at least in part with FBF-1 in repressing GLD-1 levels [21]. We choose the model proposed by Brenner and Schedl [22], where LST-1 acts in both the same pathway and in parallel to FBF-1, while SYGL-1 acts in the same pathway as FBF-1 to repress base GLD-1 levels (Figure 2A). In addition, we modeled functional redundancy of LST-1 and SYGL-1 that has been experimentally observed in *C*. *elegans* [31].

**Figure 2:**
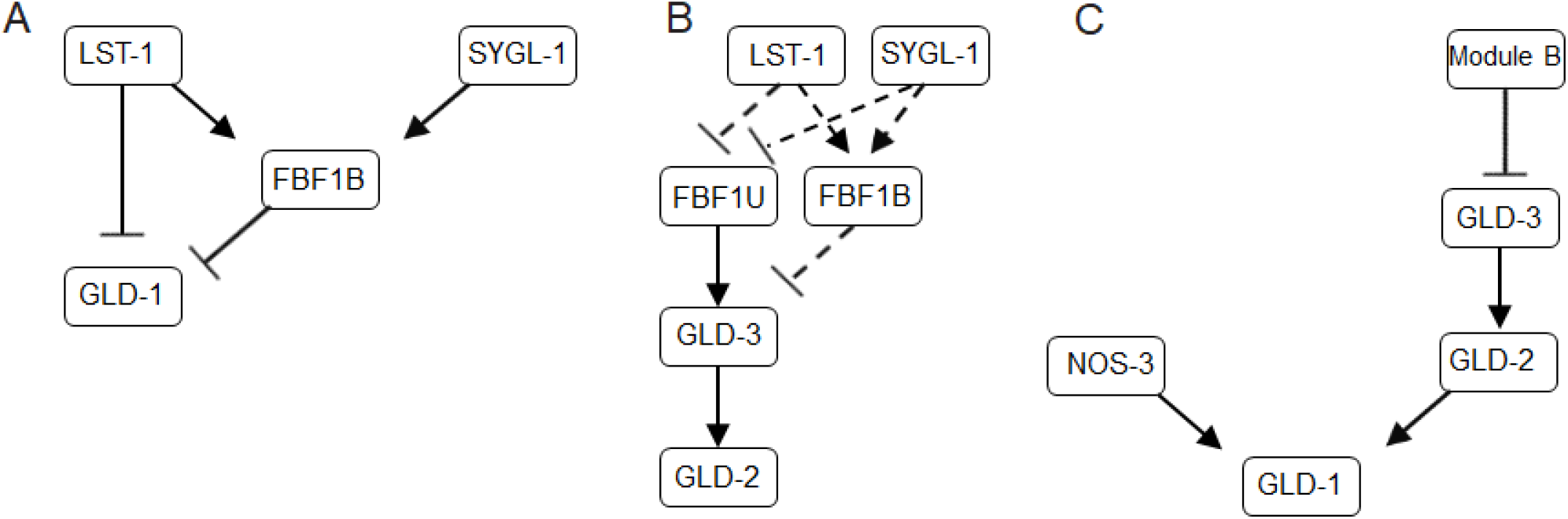
Functional modules in the gene regulatory network (A) Module A – SYGL-1 and LST-1 model. (B) Module B – two-state model of FBF-1. (C) Module C - GLD-1 accumulation model.

For the GLD-2 pathway, which includes GLD-2 and GLD-3, we constructed the module such that the dynamic behavior of FBF-1 can be further investigated. FBF- 1 was found to inhibit the GLD-2 pathway through repression of GLD-3 accumulation [35]. Conversely, FBF-1 appears to promote meiotic development through binding to the GLD2/GLD3 complex in the absence of LST-1 and SYGL-1 partners [33, 34]. This leads to the model suggesting that FBF-1 function in this module may switch in a partner-dependent manner [3].

We represent this complex partner-dependent behavior using a two-state (bound-unbound) model, where FBF-1 function is controlled by attachment and detachment of SYGL-1 or LST-1 (based on [34]). We assume two molecular states, which are required to characterize FBF-1 function with respect to SYGL-1/LST-1: a state where FBF-1 is bound to SYGL-1 or LST-1 in a molecular complex inhibiting the expression of the GLD2/GLD3 (FBF1B), and an unbound state where FBF-1 acts as a promoter of GLD-2/GLD-3 activities (FBF1U) (Figure 2B).

The GLD-1 accumulation pattern involves a number of genes (Figure 2C): (1) NOS-3, which functions upstream of GLD-1, promoting its accumulation [36] and (2) the GLD2/ GLD3 complex, which promotes the accumulation of GLD-1 only in the absence of FBF1B, SYGL-1 and LST-1 [22]. Our model includes the redundancy of NOS-3 vs GLD2/GLD3 complex as positive regulators of GLD-1 accumulation. This provides a functional module, where GLD-1 accumulation will be inhibited, as is observed in the distal end of gonad arm, unless LST-1, SYGL-1 and FBF-1 no longer repress GLD-1 levels in the absence of GLP-1 Notch activation (in the proximal part of the germ line).

The results of a recent study suggest that SCF^PROM-1^ meiotic entry pathway acts redundantly with and in parallel to the GLD-1 and/or GLD-2 pathways, and downstream of GLP-1 Notch signaling 1aling [43]. However, the genes that control the SCF^PROM-1^ activity are unknown. Hence, a simplified model to describe the interactions of these genes is that the GLP-1 signaling inhibits the meiosis-promoting activities of SCF^PROM-1^ through FBF-1. We used the same model (inhibition through FBF-1 or FBF-2) to describe the repression of meiotic chromosome axis and SC proteins, which act as additional regulators of meiotic development [38].

### 4.3. The Regulatory Network Reproduces Expected Profiles of Gene Activity

The model explores the effects-1ts of Notch signaling on the GLD-1, GLD-2 and SCF^PROM-1^ pathways that dictate the switch in fate to meiotic development. The simulated gene profiles were compared to the results of known phenotypes based on extensive experimental studies (see Table 1).

In Figure 3 the simulation results (the table of Boolean values of each gene) of a single specific network topology (see example in Supplementary Material, Figure S2) are presented. Since this is a synchronous Boolean network, if the state of all components repeats itself in two consecutive time steps, the system will remain in this state indefinitely. The simulations describe genetic interactions in an *a priori* stem cell that is experiencing Notch signaling activity by virtue of its distal position in the germ line. Hence, the DTC signal was modeled as constitutively active (by means of self-activating signal (S0), causing the core genes to be active over the course of the simulation. The expected outcome in this example after 20 steps is *StemcellFate*.

**Figure 3:**
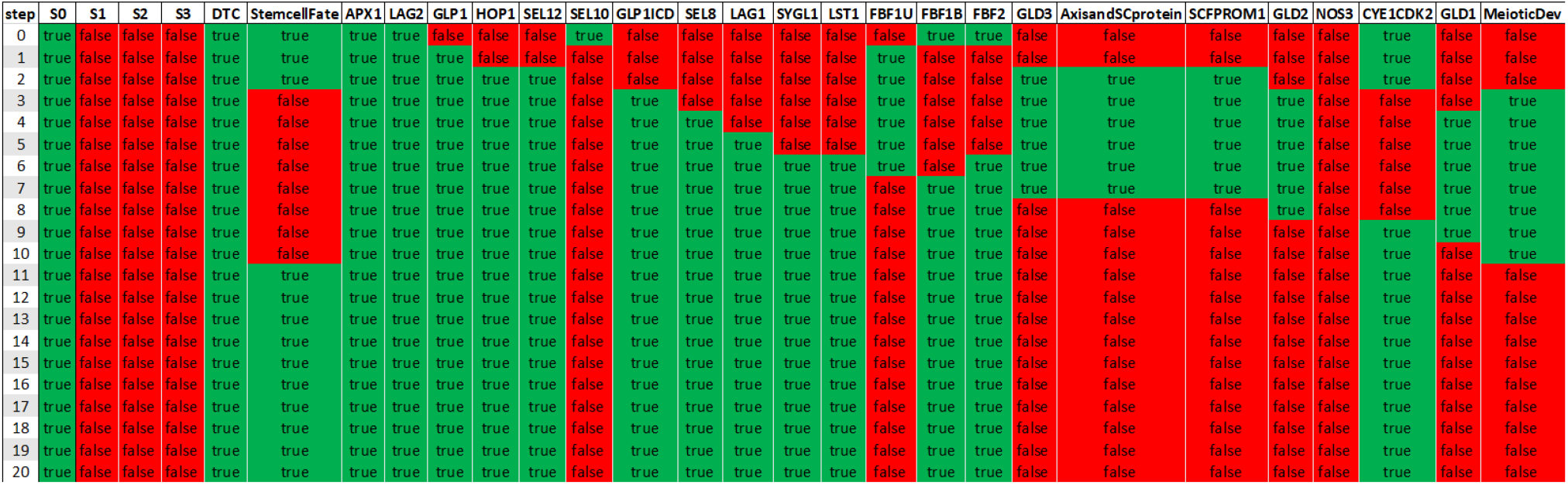
The table of Boolean values of each gene of a specific network topology. Each column corresponds to a specific *gene activity* (sorted by order of activation) throughout the simulation. Each row corresponds to the state of the genes at a different time step, with a green rectangle indicating that a gene is active, and a red rectangle indicating that a gene is inactive. The DTC signal is modeled as constitutively active, promoting the stem cell fate.

Next, we analyzed the architecture and the order of *gene activity* in two different setups (see Supplementary Material, Figure S3 and S6). One setup allows to simulate a stem cell that loses DTC signaling as it moves from the distal-most end of the germ line further proximally. Another setup represents a cell further from the DTC in the proliferative zone by reducing but not eliminating the DTC signal duration (DTC = 1 at *t* = 0 only). Once we confirmed that the literature derived regulatory network reproduces the expected gene expression profiles of young adult hermaphrodite germline stem cell system, we proceeded to conduct *in silico* perturbations.

### 4.4. Comparing Knock-out and Overexpression Simulations to Experimental Data

To test our model’s behavior under *in silico* perturbations we chose to simulate a number of conditions with known effects on the stem cell fate or meiotic development (see Table 2). The encoded experimental observations can be found in Supplementary Files. The following simulations demonstrate how a cell residing adjacent to the DTC niche responds to mutations. As mentioned, the DTC functions to promote the stem cell fate/inhibit meiotic development through GLP-1 Notch signaling. Loss of GLP-1 signaling activity causes all previously established stem cells to eventually undergo meiotic entry. GLP-1 activity can be removed during the adult stage using a conditional reduction-of-function allele of *glp-1*. When this mutant is reared at the restrictive temperature, the phenotype is indistinguishable from the *glp-1(null)* loss-of-function [40]. Therefore, we can define the GLP-1 gene as knocked out in the reasoning framework, simulating loss of GLP-1 activity (starting at time step *t* = 0) in the young adult germ line (Supplementary File - Experiment Five). The simulations show that in the absence of GLP-1 at step *t* = 0, even though DTC is constitutively active, none of the core Notch pathway genes within the scope of the model are activated, whPiRleOtMh-e1 meiotic entry pathways (the GLD-1, GLD-2 and SCF^PROM-1^) promote meiotic entry (Figure 4A). Furthermore, the absence of GLP-1 has the same effect on the cell fate decision in the case of degrading DTC signaling (not shown). The results of the model simulations are consistent with experiments in the *glp-1* mutant, where following loss of GLP-1 signaling activity all germ cells that would normally divide mitotically in the presence of GLP-1 siganling, enter meiosis [40].

**Figure 4:**
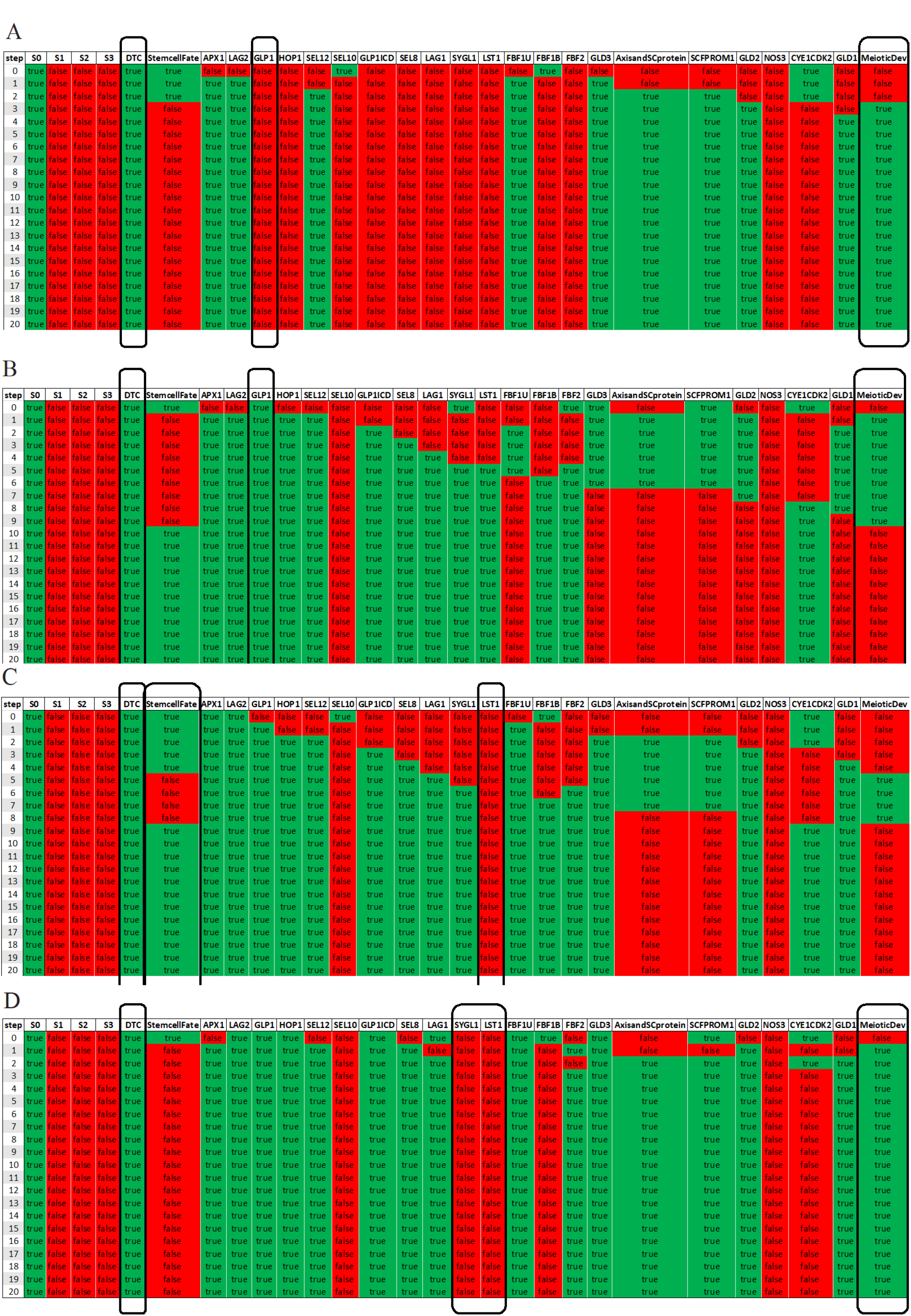
*In silico* genetic perturbations. KO and constitutive activity of GLP-1 (A,B), KO of LST-1 and LST-1 SYGL-1 (C,D).

Next, we simulated constitutive activity of GLP-1 Notch signaling by defining the *glp-1* gene as constitutively active in the reasoning framework (Supplementary File - Experiment Ten). It is known that constitutive activity of GLP-1 prevents entry into meiosis [40]. In all the solutions, meiotic development is not attained upon constitutive GLP-1 signaling, in line with investigations by Berry et al. [42], as indicated by the constant activation of the core Notch pathway genes (Figure 4B).

The effector genes *lst-1* and *sygl-1* function redundantly to promote the stem cell fate downstream of GLP-1 activity. Experiments demonstrate that either an *lst-1* or *sygl-1* single deletion mutation does not interfere with the germline stem cell fate, indicating that each is sufficient for maintaining stem cell fate [31]. In our model simulations, knock-out of either gene alone (Supplementary File - Experiment Eight and Nine) results in normal stem cell fate retention (see LST-1 knock-out example in Figure 4C) as compared to the nonperturbed network (Figure 3). In contrast, performing the *lst-1 sygl-1* double mutant (Supplementary File - Experiment Six) in the presented regulatory network model exhibited meiotic entry (Figure 4D) similar to the complete loss of GLP-1 Notch signaling (Figure 4A) and in accordance with the *in vivo* results [31].

In the setup with constitutive activity of either LST-1 or SYGL-1 (Supplementary File - Experiment Eleven and Twelve), the model exhibited failure to enter meiosis as the outcome of the simulations (not shown). *In vivo*, overexpression of LST-1 or SYGL-1 proteins in the presence of FBF-1 (a key partner of mRNA repression in stem cells) generates an overproliferation phenotype and leads to germline tumor formation [34].

The GLD-1 and GLD-2 pathways have been demonstrated to promote meiotic development together with the SCF^PROM-1^ pathway. As shown in Kadyk and Kimble [41], the germline phenotype of *gld-2(0) gld-1(0)* double mutants is a meiotic entry defect. That is, the mutant germ line is mitotic throughout, forming meiotic entry-defective tumors. The knock-out simulation of *gld-1 gld-2* combination (Supplementary File - Experiment Seven) in the presented genetic regulatory network model shows a failure to enter meiosis (Figure 5).

**Figure 5:**
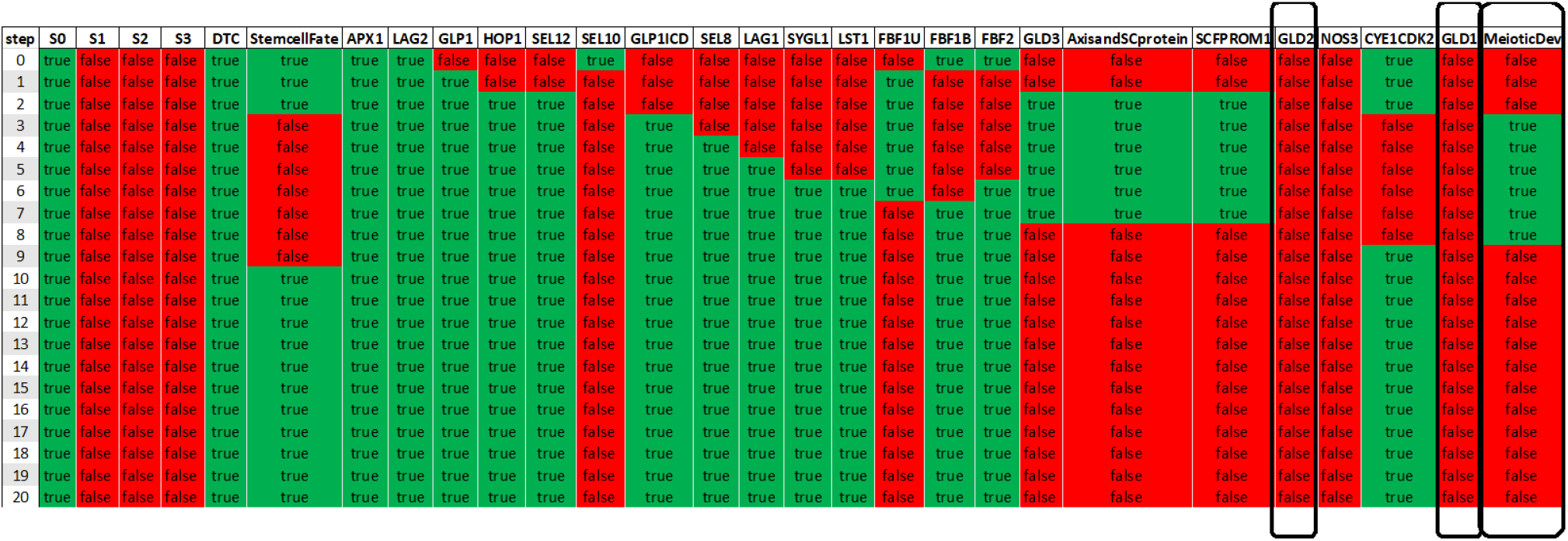
Knock-out of GLD1 GLD2 combination.

In the setup with degrading DTC signaling, the results *in silico* perturbations described in this section were similar to that obtained with the constitutively active DTC (not shown). In summary, in each case, the simulations recapitulated the *in vivo* observations.

### 4.5. Testing the Null Hypothesis for a Range of Genetic Perturbations

Our reasoning framework enables exploration of the new behavior of the presented network and finding new patterns, if any, that have not yet been experimentally observed. The solver can search for solutions that satisfy the set of defined experimental constraints based on the network topology even if they contradict experimentally observed behavior. This type of test is termed here the *null hypothesis* (the opposite of the accepted hypothesis).

For example, we previously tested whether *gld-1 gld-2* double knock-out leads to failure to enter meiosis (Figure 5). We also tested the *null hypothesis* – that the same knockout condition might result in meiotic entry. For this purpose, we use the same experimental constraint for the initial state and an opposite constraint for the final state of the simulation (see Supplementary File). The results of this simulation clearly showed that for the GLD-1 GLD-2 double knock-out the *null hypothesis* is satisfiable. Therefore, it was concluded that some models predict that the GLD-1 GLD-2 double knock-out will lead to failure to enter meiosis, while others predict that this knock-out might not lead to failure to enter meiosis. In this particular case, the result that meiotic entry occurs in the GLD-1 GLD-2 double-KO is possible due to activity from the remaining SCF^PROM-1^ pathway that continues to promote meiotic entry in the absence of both GLD-1 and GLD-2 (Figure 6).

**Figure 6:**
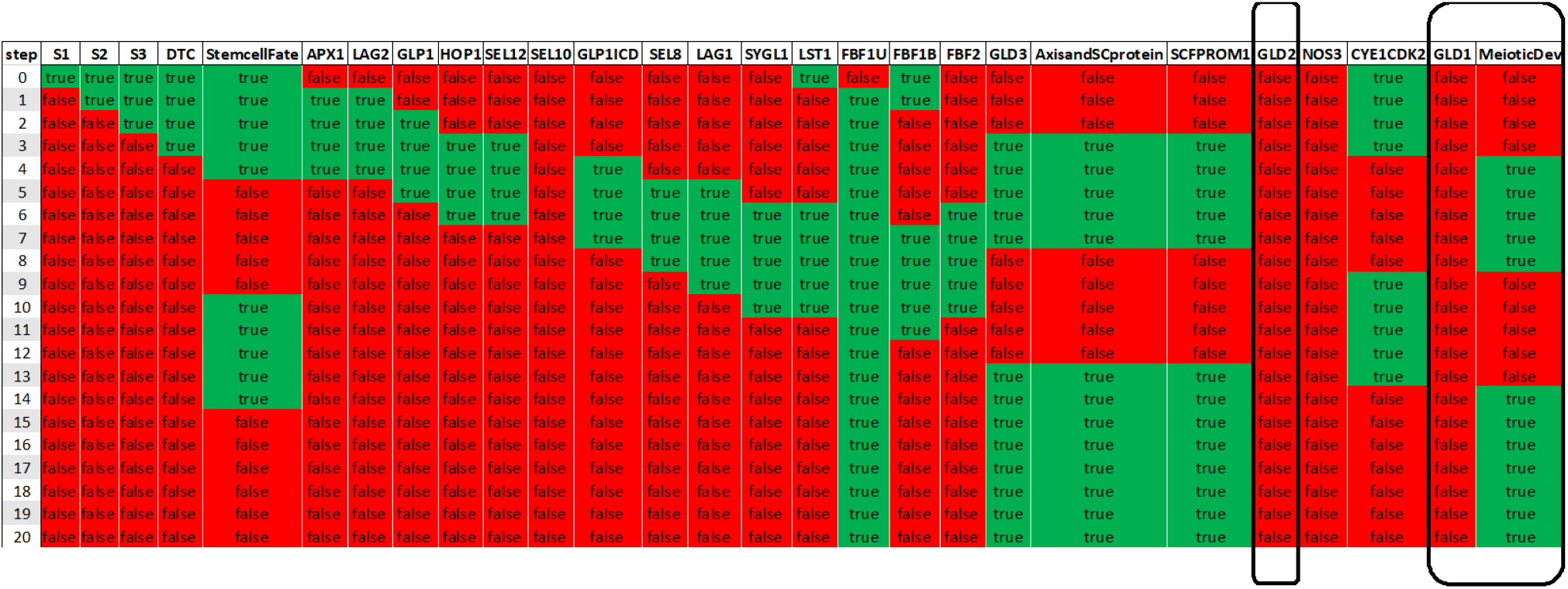
Null hypothesis for GLD1/GLD2 knock-out.

Next, we tested the *null hypothesis* for all the previously mentioned knockout and “overexpression” conditions (per Table 2). We found that for all the existing models the *null hypothesis* is unsatisfiable, which is consistent with *in vivo* studies.

## 5. Discussion

In this study, we present a new computational model representing the genetic regulatory network that controls the stem cell fate versus meiotic development decision in the young adult *C*. *elegans* hermaphrodite germ line. This model allowed us to describe how the on/off states of *gene activity* (as gleaned from both genetic and biochemical experiments reported in the experimental literature) and their interactions lead to a cell fate choice (either an undifferentiated state - “stem cell fate” or to enter into a differentiated state - “meiotic development”) and are consistent with experimental evidence.

### 5.1. The Network Model Describes Germline Stem Cell Decision

We show that simulations of the current model reproduce the overall sequence of gene states over the course of decision-making steps. As discussed below, the predictive power of our model was tested by comparing the simulated order of the gene activation/repression with previously published experimental data.

The process of developing this regulatory network highlighted possible gaps in our knowledge. For example, when we simulated a stem cell in the proliferative zone responding to reduced level of the DTC signal, only some of the simulations resulted in a meiotic development decision, implying that some other component promotes specification of the stem cell fate. This modeling result led to the identification of the need to impose regulation of the CYE1/CDK2 complex in promoting the proliferative fate; the activity of this complex was not initially regulated in our model. Indeed, when we reduce CYE1/CDK2 activity, together with a low DTC signaling activity, the network attains meiotic development decision. We modeled this by introducing an external self-degrading signal that controls CYE1/CDK2 activity from initiation of the simulation.

This example illustrates how the combination of experimental knowledge and computational model can give important clues about the role of CYE1/CDK2 in proliferative fate specification or can reveal missing regulatory interactions that control its expression. Although CYE1 and CDK2 are known as regulators of G1/S cell cycle phases, their expression in stem cells is phase-independent [39, 44], and CDK2 levels are controlled transcriptionally by glycogen synthase kinase GSK-3 through inhibition of transcription factor DPL-1 [45].

We further simulated a range of genetic perturbations (KO and FE) and compared the results with known abnormal phenotypes from several experimental studies. The order of *gene activity* matches expectations of intermediate gene profiles and ends in cell fate choice of observed phenotypes [31, 34, 40, 41, 42]. For example, constitutive activity of GLP-1 signaling activity leads to failure to enter meiosis. This simulation result is consistent with the DTC ablation in *glp-1(oz112)* animals, where germ cells never leave the mitotic cell cycle [42].

### 5.2. Formulating Model Predictions

To determine whether a genetic perturbation can result in a biological behavior that has yet to be experimentally observed, we tested the *null hypothesis* for each case. This analysis investigated whether some, all or none of the models contradict expected behavior and what is a mechanism through which this behavior is achieved. The results of these simulations show that for all the existing models, except the GLD-1 GLD-2 double KO model, the *null hypothesis* is unsatisfiable. This means that all these models predict the behavior, which is consistent with *in vivo* studies. The GLD-1 GLD-2 KO hypothesis states that *gld-1 gld-2* combination knock-out leads to failure to enter meiosis [41]. For the GLD-1 GLD-2 KO model, we found that both the KO and the *null hypothesis* are satisfiable independently. Do germ cells enter meiotic prophase normally despite the knock-out of two out of three meiotic entry pathways? This scenario was ruled out by several experimental studies [41, 43, 46]. Moreover, several studies suggest that the relative strength of each pathway is different [36, 43, 46, 47]. In our computational model, the outcome of this simulation is possible because the solver gives equal weight to each of these components (GLD-1, GLD-2, SCF^PROM-1^ and meiotic chromosome axis and SC protein). This simulation is an example of how the capability of the solver to make predictions of multiple biological behaviors is counteracted by the absence of quantitative information, which is essential for understanding of many genetic regulatory mechanisms on the molecular level.

### 5.3. Suitability of the Network Model to Simulate Various Biological Phenomena

Our network model demonstrates various important phenomena that take place in biological systems: (1) gene redundancy, (2) bound and unbound motifs, (3) dynamic behavior of extracellular/intracellular input signals, and (4) various regulatory mechanisms for individual network components. *In vivo*, the robustness of cell fate choice in the *C*. *elegans* germ line is partially attributed to the redundancy of *gene activities*. The effect of inactivating one gene can often be hidden by sufficient activity of its redundant partners. Examples of apparently redundant *gene activities* in our model include *hop-1* and *sel-12*, *lst-1* and *sygl-1*, and *nos-3* vs *gld-2* and *gld-3* (when acting as activators of GLD-1 accumulation). In these cases, in the current model we assume that the redundant genes are equally effective, and that each is sufficient for the normal pattern of germline development. For example, we tested whether either *lst-1* or *sygl-1* knock-out can affect germline stem cell fate based on experimental observations of Kershner et al. [31]. We showed that both *lst-1* or *sygl-1* knock-out simulations exhibited profiles of *gene activity* comparable to non-perturbed signaling, which suggests that gene redundancy can be accurately reproduced with the presented network model. Moreover, using the reasoning framework it is possible to specify more than two redundant genes per function. Future extensions of this tool will be to permit simulations of more expressive functional effects between redundant genes.

Protein-protein interactions correspond to binding and dissociation mechanisms, which depends on the number of binding sites, the binding affinity, the structural and functional properties of protein-protein complex, etc. The same protein can have multiple functions in the same biological system depending on its binding partners. Our model simulates this complex behavior using a two-state model. An example is FBF-1. By including bound and unbound states of FBF-1 (FBF-1B and FBF-1U) as distinct components of the network, we could describe a partner-switching behavior (from interacting with LST-1 and SYGL-1 in repression of the meiotic entry pathways to promotion of GLD-2/GLD-3 activities, resulting in activation of meiotic entry). A similar approach using the reasoning framework could be applied to study more complex pathways by considering several possible states of protein, transitions between them and temporal information regarding these transitions.

Using the reasoning framework, we study three possible states of the single cell: within the range of the DTC signal, out of range of the DTC signal or losing the DTC signal. These model state setups differ in the strength (duration of activation) of signaling from the DTC (by employing a constant external signal S0, without it or with three external signals S1, S2, S3). The use of “external signals” allows us to reproduce the profiles of *gene activity* similar to *in vivo* responses of Notch signaling. Depending on the specific research question or hypothesis, we can use the reasoning framework to model input signals with various dynamic behavior. For example, when simulating sustained expression of a gene, the input signal should be specified as sustained (active) throughout the experiment. Another example is loss of signaling activity, which can be modeled with a self-degrading signal (that is specified to become active for a single time step). An additional possible case is when genes show an oscillating expression pattern due to alternating protein levels or crosstalk with other pathways.

Changes of external or internal cell conditions often induce associated changes in the expression of underlying genes. Since our model is represented as a Boolean network, the state of each component (either on or off) is calculated from the state of adjacent components. The synchronous updates are used in order to avoid scenarios in which not all genes are updated throughout the experiment or certain genes are repeatedly updated, but not others. Investigating an asynchronous update scheme for this model is a topic for future work with more constrained and complex networks.

The reasoning framework uses a set of regulation rules where none, some or all component regulators (activators/repressors) are present, considering two regulators of each type. For modeling gene redundancy, the downstream components can use conditions where only one of two regulators (activators or repressors) is present. The result from applying these regulation conditions in our model is a reliable and robust demonstration of a biological function of components in the genetic network, using few activators and/or repressors as happens *in vivo*.

### 5.4. Future Extensions

In addition to the possible extensions of the reasoning framework mentioned above, our network model could be extended and refined in different ways. For example, currently the model describes the dynamics of the network that controls cell fate decisions at a single cell level. The next step towards further development of the model is to extend it to a population of cells. The combination of a detailed description of the interactions within the cell, the interplay between cells in population and their communication with the DTC niche could provide new insights into the regulation of stem cell systems. Furthermore, a detailed *in silico* implementation of a single cell provides the possibility to capture more realistic whole tissue simulations. This will open up the possibility to computationally test the relative contribution of interactions within the cell that are usually overlooked in experimentally measured phenotypes of the entire germ line.

Finally, the extended model could be used to explore motif-based design patterns and their role in controlling cellular behaviors. Simple regulatory motifs have been identified as the basic building blocks of many biological networks [48] where their structure may determine a broad range of functions (e.g., sign-sensitive delay, pulse generation). By employing formal methods for identification of these motifs and their effect on the cellular functions, one can ensure that a comprehensive analysis was done for the network of interest [49].

## Data Availability

Supplementary Files related to this article can be found at https://github.com/kuglerh/Germline. The Reasoning Engine framework is available on GitHub (https://github.com/fsprojects/ReasoningEngine).

## Acknowledgements

This work was supported by the Horizon 2020 research and innovation programme for the Bio4Comp project under grant agreement No. 732482 and by the ISRAEL SCIENCE FOUNDATION (grant No.190/19) to HK.

## Supplementary Material

### 1. Regulation Conditions

The set of 18 regulation conditions (excluding the two threshold rules) is taken from Yordanov et al. [11]. Permission to be granted. Each column represents a different condition of a target node where a specific rule is used. The state of each target (shown as a red circle in the header diagram) is updated depending on the states of its regulators (activators or repressors). Black circles indicate whether all, some or none of the regulators are active, representing several specific conditions where a target is non-inducible and non-repressible (a), only inducible (b), only repressible (c), or both inducible and repressible (d). The conditions highlighted with yellow boxes represent non-inducible and non-repressible targets that cannot be constantly activated or repressed regardless of the state of their activators.

### 2. Example of the network topology

In Figure S2 below, an example of the network topology of a single specific solution is presented. The positive regulations appear as solid black lines, while red color of the arrows corresponds to the negative regulations. The *optional* interactions instantiated by the solver in this solution together with assigned regulation conditions satisfy all observations, showing how the expected behavior can be realized in this model (for details on how regulations conditions are used to determine the state of the regulated component, see [11] and Figure S1).

### 3. Individual cell’s setup

This setup allows to simulate a stem cell that loses DTC signaling as it moves from the distal-most end of the germ line further proximally. Therefore, the specified initial state of the simulation along with DTC signal is *StemcellFate* and the expected outcome in this example after 20 steps is meiotic development *MeioticDev*. To model degrading DTC signaling (from present to absent) inputs were separated into 3 generic signals that we call signal 1, 2, and 3 (S1, S2, S3) and that are connected in series (see Figure S1) resulting in the absence of DTC signal from *t* = 4 onwards. The initiation of the network interactions starts when the DTC signals to the germ line by expressing LAG-2 and APX-1 proteins [23], [24] at time step *t* = 1. APX-1 and LAG-2 redundantly activate GLP-1 Notch receptor, which results in the release of the GLP-1 intracellular domain [25]. Other GLP-1 signaling pathway genes *hop-1* and *sel-12* are activated at *t* = 3 of the simulation. The network state at *t* = 4 captures the formation of a ternary complex containing GLP1ICD, LAG-1, and SEL-8 [26] (see Figure S2). At the next steps, GLP-1 signaling leads to the activation of *lst-1* and *sygl-1* genes and thereby to the inhibition of the GLD-1, GLD-2 and SCF^PROM-1^ meiotic entry pathways through FBF. Once FBF is inactivated, meiotic entry pathways return to being active and *MeioticDev* is constituted from time step *t* = 15.

Finally, we simulated a cell further from the DTC in the proliferative zone by reducing but not eliminating the DTC signal duration (DTC = 1 at *t* = 0 only). The analysis of these simulations showed that a reduction of the DTC signal is not enough to ensure that the cell always undergoes meiotic development. While in some simulations the cell enters meiotic development state (Figure S4), in others it exhibits transient recurrence of stem cell fate (Figure S5).

Next, we checked whether some network component promotes stem cell fate, despite a reduced level of DTC signaling. We found that CYE1/CDK2 (cyclin E and cyclin-dependent kinase) appears to be an active regulator of the stem cell fate. Therefore, we specified external selfdegrading signal, which reduces CYE1/CDK2 activity for realistic simulation of this scenario (Figure S6). This is consistent with investigation by Fox et al. [39], which showed that CYE1/CDK2 plays important role in promoting the proliferative fate, specifically in cells with reduced GLP-1 signaling.

---

**Figure S1.**
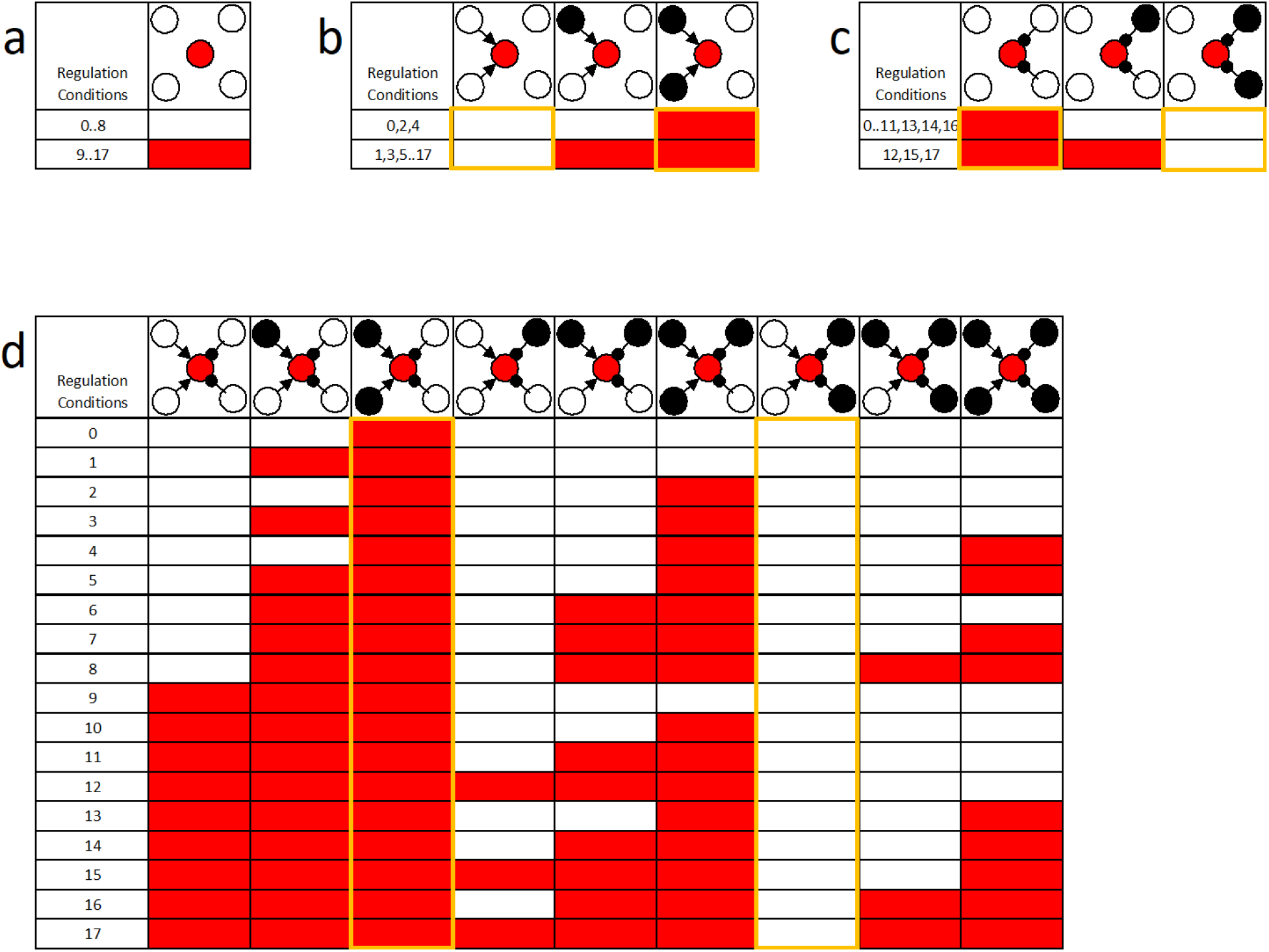

**Figure S2.**
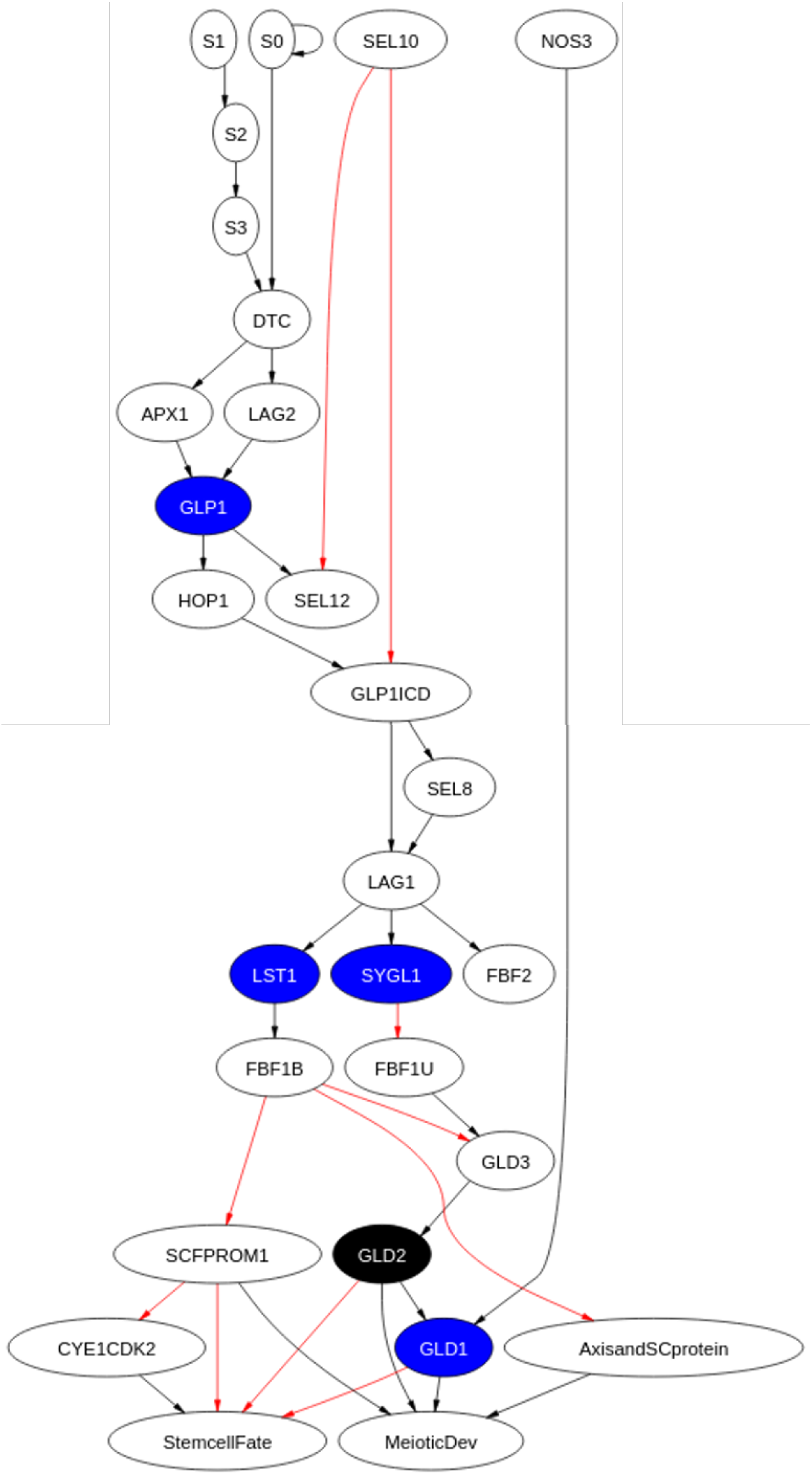

**Figure S3.**
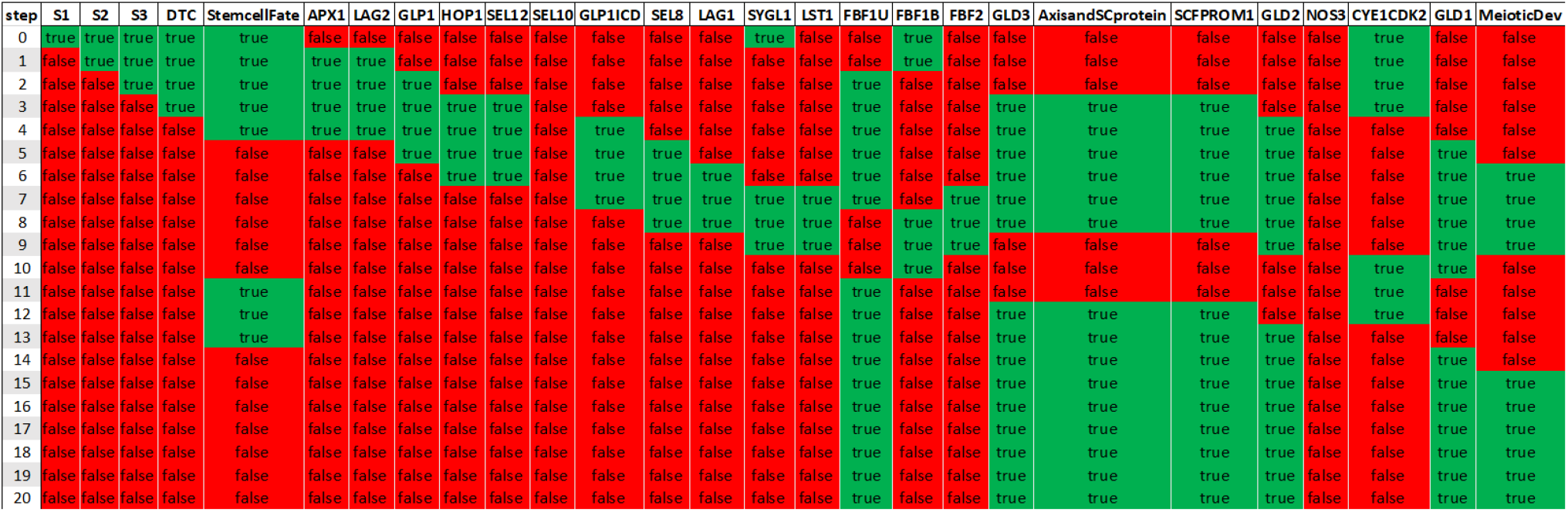

**Figure S4.**
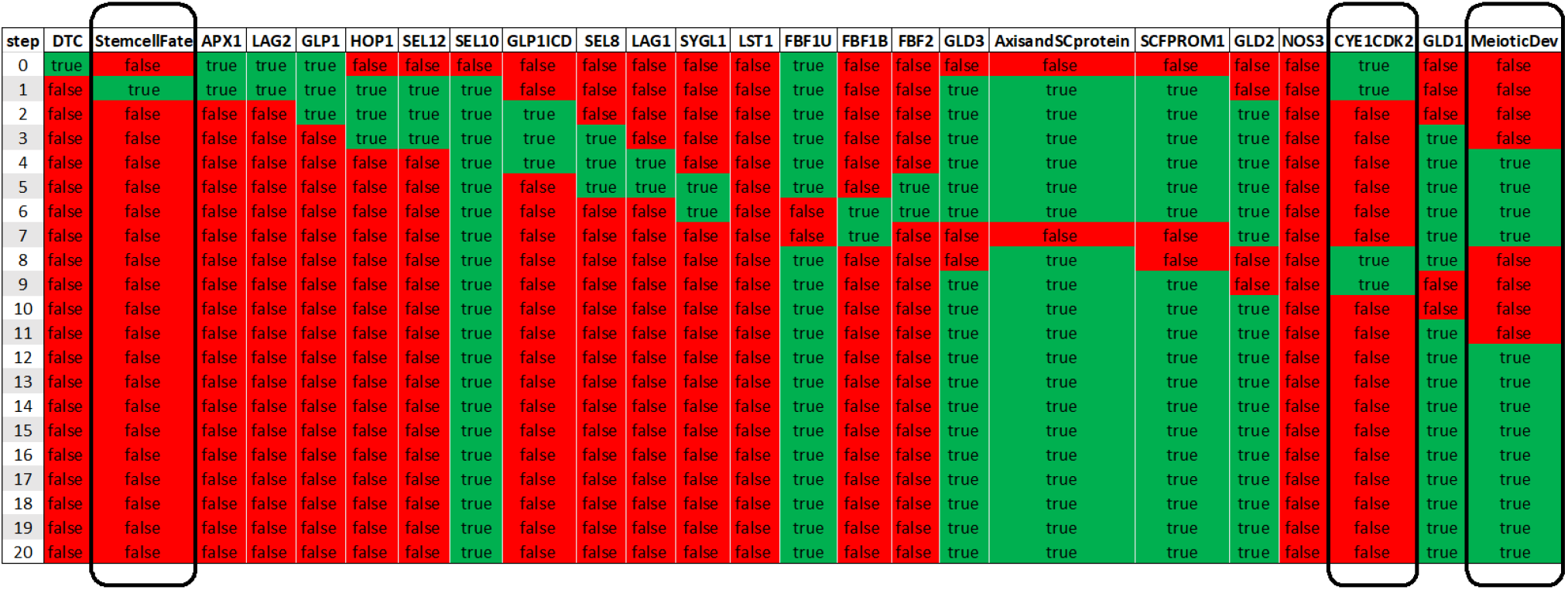

**Figure S5.**
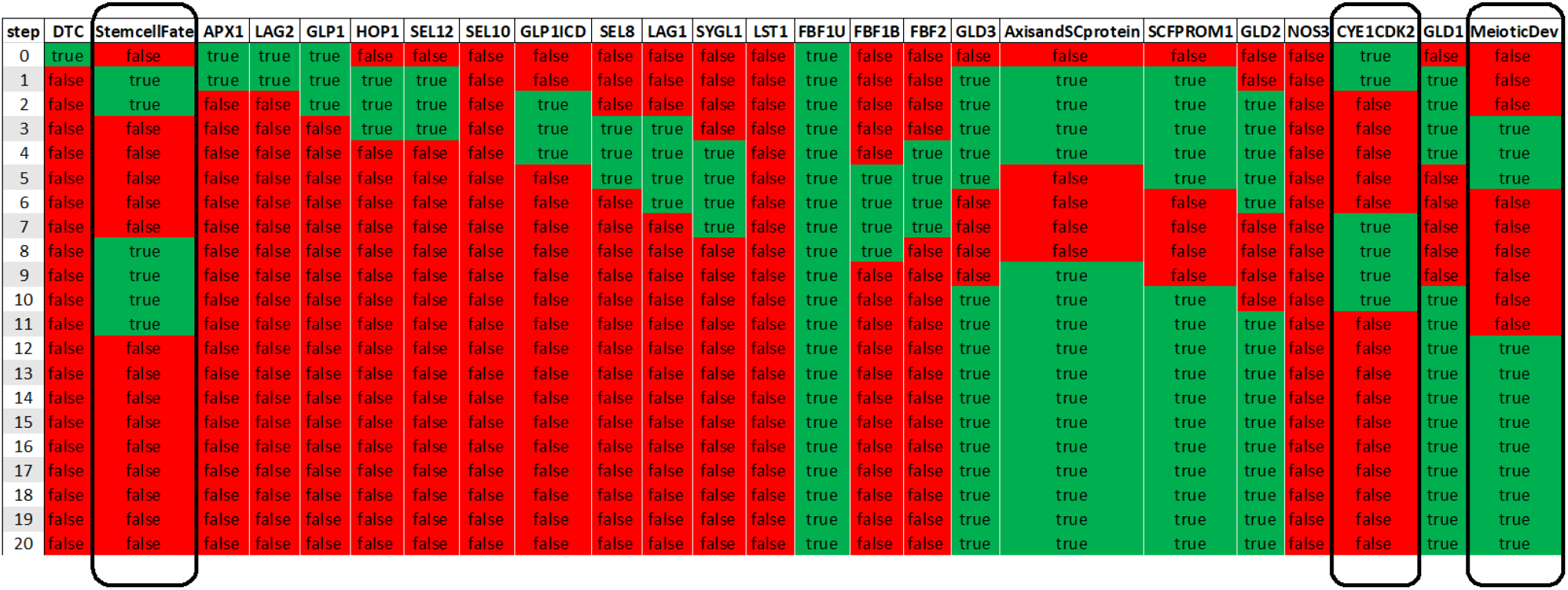

**Figure S6.**
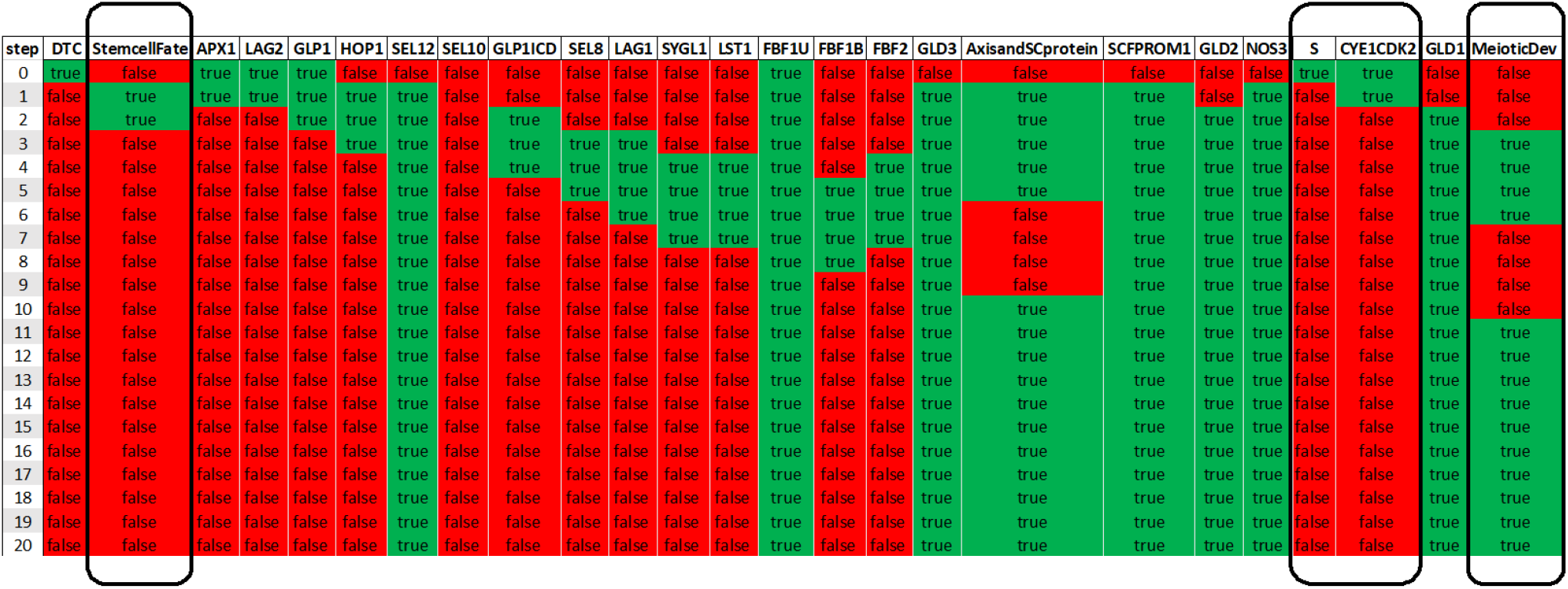

## References

[1] Simons, B.D., Clevers, H.: Strategies for Homeostatic Stem Cell Self-Renewal in Adult Tissues. Cell 145(6), 851–862 (2011)

[2] Batlle, E., Clevers, H.: Cancer stem cells revisited. Nature Medicine 23, 1124–1134 (2017)

[3] Hubbard, E.J.A., Schedl T.: Biology of the Caenorhabditis elegans Germline Stem Cell System. Genetics 213(4), 1145–1188 (2019)

[4] Atwell, K., Dunn, S.-J., Osborne, J.M., Kugler, H., Hubbard, E.J.A.: How computational models contribute to our understanding of the germ line. Molecular reproduction and development 83(11), 944–957 (2016)

[5] Setty, Y., Dalf′o, D., Korta, D., Hubbard, E.J.A., Kugler, H.: A model of stem cell population dynamics: in silico analysis and in vivo validation. Development 139(1), 47–56 (2012)

[6] Atwell, K., Qin, Z., Gavaghan, D., Kugler, H., Hubbard, E.J.A., Osborne, J.M.: Mechano-logical model of C. elegans germ line suggests feedback on the cell cycle. Development 142(22), 3902–3911 (2015)

[7] Harel, D.: Statecharts: A visual formalism for complex systems. Science of Computer Programming 8(3), 231–274 (1987)

[8] Harel, D., Kugler, H.: The Rhapsody semantics of statecharts (or, on the executable core of the UML. Integration of Software Specification Techniques for Applications in Engineering, 325–354, Springer, Berlin, Heidelberg (2004)

[9] Dunn, S.-J., Martello, G., Yordanov, B., Emmott, S., Smith, A.G.: Defining an essential transcription factor program for naive pluripotency. Science 344(6188), 1156–1160 (2014)

[10] Goldfeder, J., Kugler, H.: BRE:IN - A Backend for Reasoning About Interaction Networks with Temporal Logic. Computational Methods in Systems Biology 11773, 289–295 (2019)

[11] Yordanov, B.,Dunn, S.-J., Kugler, H., Smith, A., Martello, G., Emmott, S.: A method to identify and analyze biological programs through automated reasoning. NPJ Systems Biology and Applications 2(16010) (2016)

[12] Guziolowski, C., Videla, S., Eduati, F., Thiele, S., Cokelaer, T., Siegel, A., Saez-Rodriguez, J.: Exhaustively characterizing feasible logic models of a signaling network using answer set programming. Bioinformatics 29(18), 2320–2326 (2013)

[13] Koksal, A., Pu, Y., Srivastava, S., Bodik, R., Fisher, J., Piterman, N.: Synthesis of biological models from mutation experiments. In: SIGPLAN-SIGACT symposium on principles of programming languages. ACM (2013)

[14] Sharan, R., Karp, R.M.: Reconstructing boolean models of signaling. Journal of Computational Biology 20(3), 249–257 (2013)

[15] Rosenblueth, D.A., Muñoz, S., Carrillo, M., Azpeitia, E.: Inference of boolean networks from gene interaction graphs using a sat solver. In: International Conference on Algorithms for Computational Biology. pp. 235–246. Springer (2014)

[16] Koksal, A.: Program Synthesis for Systems Biology. Ph.D. thesis, University of California at Berkeley (2018), technical Report No. UCB/EECS-2018-49

[17] Chevalier, S., Froidevaux, C., Paulevé, L., Zinovyev, A.: Synthesis of boolean networks from biological dynamical constraints using answer-set programming. In: 2019 IEEE 31st International Conference on Tools with Artificial Intelligence (ICTAI). pp. 34–41 (2019). 10.1109/ICTAI.2019.00014

[18] Razzaq, M., Kaminski, R., Romero, J., Schaub, T., Bourdon, J., Guziolowski, C.: Computing diverse boolean networks from phosphoproteomic time series data. In: International Conference on Computational Methods in Systems Biology. pp. 59–74. Springer (2018)

[19] Chevalier, S., Noël, V., Calzone, L., Zinovyev, A., Paulevé, L.: Synthesis and simulation of ensembles of boolean networks for cell fate decision. In: Abate, A., Petrov, T., Wolf, V. (eds.) Computational Methods in Systems Biology. pp. 193–209. Springer International Publishing, Cham (2020)

[20] Biane, C., Delaplace, F.: Causal Reasoning on Boolean Control Networks Based on Abduction: Theory and Application to Cancer Drug Discovery. IEEE/ACM Trans. Comput. Biol. Bioinformatics 16(5), 1574–1585. (2019)

[21] Chen, J., Mohammad, A., Pazdernik, N., Huang, H., Bowman, B., Tyckse, E., Schedl, T.: GLP-1 Notch—LAG-1 CSL control of the germline stem cell fate is mediated by transcriptional targets lst-1 and sygl-1. PLoS Genetics 16(3), e1008650 (2020)

[22] Brenner, J.L., Schedl, T.: Germline stem cell differentiation entails regional control of cell fate regulator GLD-1 in Caenorhabditis elegans. Genetics 202(3), 1085–1103 (2016)

[23] Nadarajan, S., Govindan, J.A., McGovern, M., Hubbard, E.J.A., Greenstein, D.: MSP and GLP1/Notch signaling coordinately regulate actomyosin-dependent cytoplasmic streaming and oocyte growth in C. elegans. Development 136(13), 2223–2234 (2009)

[24] Henderson, S.T., Gao, D., Lambie, E.J., Kimble, J.: lag-2 may encode a signaling ligand for the GLP-1 and LIN-12 receptors of C. elegans. Development 120(10), 2913–2924 (1994)

[25] Greenwald, I., Kovall, R.A.: Notch signaling: genetics and structure. WormBook, ed. The C. elegans Research Community, 1–28 (2013)

[26] Petcherski, A.G., Kimble, J.: LAG-3 is a putative transcriptional activator in the C. elegans Notch pathway. Nature 405, 364–368 (2000)

[27] Agarwal, I., Farnow, C., Jiang, J., Kim, K.-S., Leet, D.E., Solomon, R.Z., Hale, V.A., Goutte, C.: HOP-1 Presenilin Deficiency Causes a Late-Onset Notch Signaling Phenotype That Affects Adult Germline Function in Caenorhabditis elegans. Genetics 208(2), 745–762 (2018)

[28] Levitan, D., Greenwald, I.: Facilitation of lin-12-mediated signalling by sel-12, a Caenorhabditis elegans S182 Alzheimer’s disease gene. Nature 377, 351–354 (1995)

[29] Hubbard, E.J.A., Wu, G., Kitajewski, J.K., Greenwald, I.: sel-10, a negative regulator of lin-12 activity in Caenorhabditis elegans, encodes a member of the CDC4 family of proteins. Genes and Development 11(23), 3182–3193 (1997)

[30] Wu, G., Hubbard, E.J.A., Kitajewski, Greenwald, J.K.: Evidence for functional and physical association between Caenorhabditis elegans SEL-10, a Cdc4p-related protein, and SEL-12 presenilin. Proc. Nat’l Academy of Sciences USA 95(26), 15787–15791 (1998)

[31] Kershner, A.M., Shin, H., Hansen, T.J., Kimble, J.: Discovery of two GLP-1/Notch target genes that account for the role of GLP-1/Notch signaling in stem cell maintenance. Proc. Nat’l Academy of Sciences USA 111(10), 3739–3744 (2014)

[32] Lamont, L.B., Crittenden, S.L., Bernstein, D., Wickens, M., Kimble, J.: FBF-1 and FBF-2 regulate the size of the mitotic region in the C. elegans germline. Developmental Cell 7(5), 697–707 (2004)

[33] Hansen, D., Schedl, T.: The regulatory network controlling the proliferation-meiotic entry decision in the Caenorhabditis elegans germ line. Current Topics in Developmental Biology 76, 185–215 (2006)

[34] Shin, H., Haupt, K.A., Kershner, A.M., Kroll-Conner, P., Wickens, M., Kimble, J.: SYGL-1 and LST-1 link niche signaling to PUF RNA repression for stem cell maintenance in Caenorhabditis elegans. PLoS Genetics 13(12), e1007121 (2017)

[35] Eckmann, C., Crittenden, S.L., Suh, N., Kimble, J.: GLD-3 and control of the mitosis/meiosis decision in the germline of Caenorhabditis elegans. Genetics 168(1), 147–160 (2004)

[36] Hansen, D., Wilson-Berry, L., Dang, T., Schedl, T.: Control of the proliferation versus meiotic development decision in the C. elegans germline through regulation of GLD-1 protein accumulation. Development 131(1), 93–104 (2004)

[37] Jantsch, V., Tang, L., Pasierbek, P., Penkner, A., Nayak, S., Baudrimont, A., Schedl, T., Gartner, A., Loidl, J.: Caenorhabditis elegans prom-1 is required for meiotic prophase progression and homologous chromosome pairing. Molecular biology of the cell 18(12), 4911–4920 (2007)

[38] Merritt, C., Seydoux, G.: The Puf RNA-binding proteins FBF-1 and FBF-2 inhibit the expression of synaptonemal complex proteins in germline stem cells. Development 137(11), 1787–1798 (2010)

[39] Fox, P.M., Vought, V.E., Hanazawa, M., Lee, M.H., Maine, E.M., Schedl, T.: Cyclin E and CDK-2 regulate proliferative cell fate and cell cycle progression in the C. legans germline. Development 138(11), 2223–2234 (2011)

[40] Austin, J., Kimble, J.: glp-1 is required in the germ line for regulation of the decision between mitosis and meiosis in C. elegans. Cell 51(4), 589–599 (1987)

[41] Kadyk, L.C., Kimble, J.: Genetic regulation of entry into meiosis in Caenorhabditis elegans. Development 125(10), 1803–1813 (1998)

[42] Berry, L., Westlund, B., Schedl, T.: Germ-line tumor formation caused by activation of glp-1, a Caenorhabditis elegans member of the Notch family of receptors. Development 124(4), 925–936 (1997)

[43] Mohammad, A., Vanden Broek, K., Wang, C., Daryabeigi, A., Jantsch, V., Hansen, D., Schedl, T.: Initiation of Meiotic Development Is Controlled by Three Post-transcriptional Pathways in Caenorhabditis elegans. Genetics 209(4), 1197–1224 (2018)

[44] White, J., Dalton, S.: Cell cycle control of embryonic stem cells. Stem Cell Reviews 1, 131–138 (2005).

[45] Furuta, T., Joo, H.-J., Trimmer, K.A., Chen, S.-Y., Arur, S.: GSK-3 promotes S-phase entry and progression in C. elegans germline stem cells to maintain tissue output. Development 145(dev161042), 1–11 (2018).

[46] Hansen, D., Hubbard, E.J.A., Schedl, T.: Multi-pathway control of the proliferation vs. meiotic development decision in the Caenorhabditis elegans germline. Developmental Biology 268(2), 342–357 (2004).

[47] Wang, L., Eckmann, C., Kadyk, L., Wickens, M., Kimble, J.: A regulatory cytoplasmic poly(A) polymerase in Caenorhabditis elegans. Nature 419, 312–316 (2002).

[48] Milo, R., Shen-Orr, S., Itzkovitz, S., Kashtan, N., Chklovskii, D., Alon, U.: Network motifs: simple building blocks of complex networks. Science 298(5594), 824–827 (2002).

[49] Kugler, H., Dunn, S.-J., Yordanov, B.: Formal analysis of network motifs. Ceška, M., Šafránek, D. (eds.) Computational Methods in Systems Biology, LNCS, vol. 11095, pp. 111–128, Cham: Springer, (2018).

[50] Kauffman, S., 1969. Metabolic stability and epigenesis in randomly constructed genetic nets. J. of Theoretical Biology 22, 437—-467.

[51] Shavit, Y., Yordanov, B., Dunn, S.-J., Wintersteiger C.M., Otani, T., Hamadi, Y., Livesey, F.J., Kugler, H.: Automated Synthesis and Analysis of Switching Gene Regulatory Networks. Biosystems 146, 26–34 (2016)

[52] Bartocci, E., Lio, P.: Computational modeling, formal analysis, and tools for systems biology. PLoS Computational Biology 12(1), e1004591 (2016)

[53] Paulevé, L., Kolčák, J., Chatain, T., Haar, S.: Reconciling qualitative, abstract, and scalable modeling of biological networks. Nature communications 11(1), 1—-7. (2020)

